# B1a B cells require autophagy for metabolic homeostasis and self-renewal

**DOI:** 10.1101/240523

**Authors:** Alexander J Clarke, Thomas Riffelmacher, Daniel Braas, Richard J Cornall, Anna Katharina Simon

## Abstract

Specific metabolic programs are activated by immune cells to fulfil their functional roles, which include adaptations to their microenvironment. B1 B cells are tissue-resident, innate-like B cells. They have many distinct properties, such as the capacity to self-renew and the ability to rapidly respond to a limited repertoire of epitopes. The metabolic pathways that support these functions are unknown. We show that B1 B cells are bioenergetically more active than B2 B cells, with higher rates of glycolysis and oxidative phosphorylation, and depend on glycolysis. They acquire exogenous fatty acids, and store lipids in droplet form. Autophagy is differentially activated in B1a B cells, and deletion of the autophagy gene *Atg7* leads to a selective loss of B1a B cells due to a failure of self-renewal. Autophagy-deficient B1a B cells downregulate critical metabolic genes and accumulate dysfunctional mitochondria. B1 B cells therefore, have evolved a distinct metabolism adapted to their residence and specific functional properties.

---

**List of non-standard abbreviations**
2-DG: 2-deoxyglucose
BCR: B-cell receptor
HSC: Haematopoietic stem cell
MMP: Mitochondrial membrane potential
NADPH: Nicotinamide adenine dinucleotide phosphate
OXPHOS: Oxidative phosphorylation
PCA: Principal component analysis
PPP: Pentose phosphate pathway
TCA: Tricarboxylic acid cycle
TMRM: Tetramethylrhodamine

## Introduction

B1 B cells are a distinct lineage of tissue-resident, innate-like B cells with critical roles in the immune response to pathogens with repetitive carbohydrate epitopes such as *S. pneumoniae* (Baumgarth, 2010). They are a major source of natural IgM, which in addition to its antimicrobial properties helps maintain tissue homeostasis by cross reaction with epitopes expressed on dead and dying cells (Chen et al., 2009). They are also an important component of barrier immunity, as they preferentially class-switch to IgA to control microbes at mucosal surfaces (Kaminski and Stavnezer, 2006). B1 B cells are normally resident in the peritoneum and pleura, although they also recirculate through secondary lymphoid tissues (Ansel et al., 2002). Following activation, they transit to the spleen or draining lymph nodes, where they secrete antibodies (Yang et al., 2007). These responses are typically antigen non-specific, as B1 B cells preferentially respond to Toll-like receptor rather than B-cell receptor (BCR) signalling (Baumgarth, 2010).

B1 B cells develop distinct from B2 cells (which include follicular and marginal zone B cells), and their developmental origins have been the subject of considerable debate (Montecino-Rodriguez and Dorshkind, 2012). B1 B cells are initially seeded following generation during fetal and early neonatal life, and the major population thereafter is maintained by self-renewal (Hayakawa et al., 1986; Krop et al., 1996). B2 B cells however, are continuously produced in the bone marrow from haematopoietic stem cells (HSCs) throughout life, although there remains limited potential for B1 production from bone marrow B1 progenitors (Barber et al., 2011). B1 B cells selection is enhanced by strong BCR signalling, which may be spontaneous or induced by self-antigens, and it has been proposed that this leads to their formation from a progenitor in common with B2 cells (the ‘selection’ model). The alternative, ‘lineage’ theory, is that B1 cells arise from a distinct progenitor (Tung et al., 2006). B1 B cells are recognised as CD19^hi^B220^lo^IgM^hi^CD23^−^; the major B1a subset is CD5^+^, the minor B1b subset CD5^−^. B1b B cells recognise a broader range of antigens, and can form memory B cells (Baumgarth, 2010).

It has become established that T lymphocytes adopt distinct metabolic programs which are highly regulated between functional subsets. Naïve T cells mainly generate energy by mitochondrial oxidative phosphorylation (OXPHOS). Upon activation, T cells additionally upregulate aerobic glycolysis, that is, reduction of pyruvate produced by glycolysis to lactate (Buck et al., 2015). OXPHOS is then downregulated as the T cell becomes a fully differentiated effector. Regulatory T cells, in comparison, predominantly generate energy by fatty acid oxidation (Michalek et al., 2011), as do memory T cells, which is thought to reflect their residence in lipid rich microenvironments such as the skin, lymph node, and intestinal lamina propria (Pearce et al., 2009; Pan et al., 2017). Innate lymphoid cells have also recently been shown to predominantly utilise environmental fatty acids (Wilhelm et al., 2016).

In contrast, comparatively little is known about the metabolic phenotypes of non-malignant B cells, and in particular the metabolic programs which maintain B cell homeostasis *in vivo* have been much less explored (Pearce and Pearce, 2013). The distinct tissue residence of B1a B cells in the peritoneum, which is a highly lipid rich environment, coupled with their self-renewal capacity and state of pre-activation suggests that they may have evolved a specific metabolic program to support these characteristics. Importantly, chronic lymphoid leukaemia is thought to frequently arise from B1 B cells, and therefore understanding their underlying metabolism may lead to new therapeutic insights (Montecino-Rodriguez and Dorshkind, 2012).

Here, we show that B1a B cells engage a metabolic program distinct from follicular B2 B cells (Fo B2). They have active glycolysis and fatty acid synthesis, with little metabolic flexibility. They acquire exogenous lipids and maintain intracellular fat stores. They are dependent, unlike Fo B2 B cells, on autophagy to survive and self-renew, and loss of autophagy causes global metabolic dysfunction and failure of lipid and mitochondrial homeostasis.

## Results and discussion

### B1a B cells have a distinct metabolic gene transcription identity

To determine if differences exist in the expression of key metabolic genes between CD5^+^CD23^−^ peritoneal B1a and splenic CD23^+^ follicular B2 B cells (Fo B2), we performed multiplex qRT-PCR using Fluidigm Biomark on cells sorted by flow cytometry (Figure 1A). Principal component analysis revealed clear separation between the cell types, as did unsupervised hierarchical clustering (Figure 1B-C). We found significantly higher expression of genes critical for glucose uptake and commitment to glycolysis (*Slc2a1* [Glut1], *Hk2* [Hexokinase 2], *Myc* [c-Myc]), regulation of fatty acid synthesis (*Acacb* [Acetyl-CoA carboxylase 2]), and lipid droplet formation (*Plin3* [Perilipin-3]) in B1a B cells compared with Fo B2 B cells (Figure 1D). Post-hoc analysis of other fatty acid metabolic genes also showed upregulation of *Acly* (ATP citrate lyase – which catalyses the formation of acetyl-CoA from citrate for fatty acid synthesis) (Figure 1D). Gene expression data were therefore suggestive of a distinct metabolic program in B1a B cells compared with Fo B2 cells, characterised by high expression of glycolysis genes, but also those important in fatty acid metabolism, synthesis, and storage.

**1.**
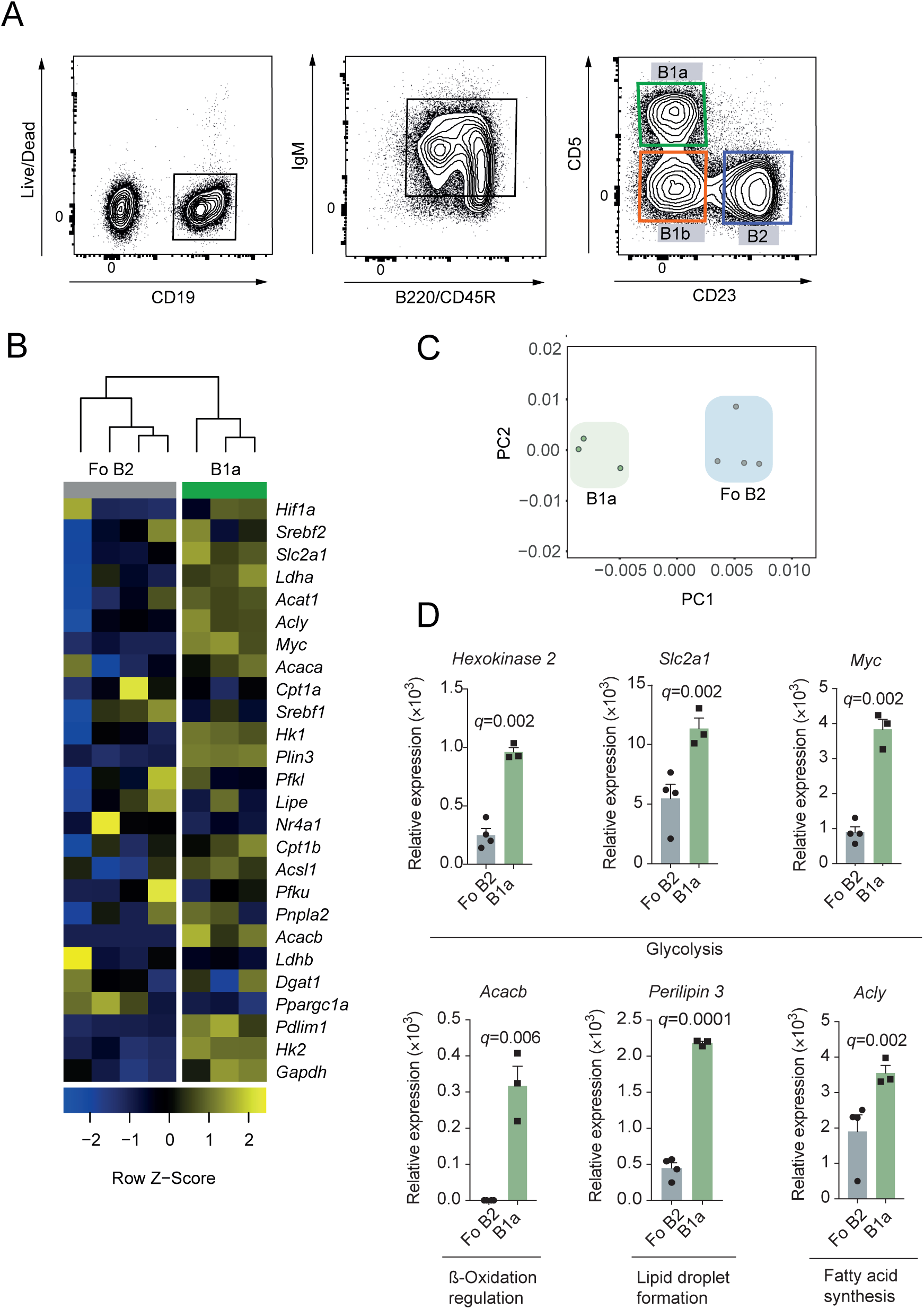
B1a B cells have a distinct metabolic gene transcription identity. **A**. Gating strategy for the identification of murine peritoneal B cell subsets. B1a B cells were identified as viable, CD19^+^, IgM^hi^, B220^lo^, CD23^−^, CD5^+^. B2 B cells were CD23^+^CD5^−^. B1b B cells were CD23^−^CD5^−^. **B**. Heatmap of Fluidigm Biomark qRT-PCR gene expression data for peritoneal B1a B cells (CD19^+^C23^−^CD5^+^) and follicular splenic B2 B cells (CD19^+^CD23^+^). One thousand cells were sorted by flow cytometry according to the gating strategy in (A), and qRT-PCR performed for a curated metabolic gene set (Supplementary Table 1). Data are target gene expression relative to β-actin, and heat map colouring is based on row Z-score. Each data point represents one C57BL/6 wild-type mouse, and is the mean of two technical replicates. Hierarchical clustering is unsupervised. **C**. Principal component analysis of data from **B**. First and second principal components are plotted. **D**. Gene expression data from **B**. Each data point is one mouse, with two technical replicates each. Mean ±SEM is depicted. Unpaired *t*-test *p*-value is adjusted for multiple testing using FDR method (5% threshold). The post-hoc *p*-value for *Acly* is presented unadjusted. Each significantly differentially expressed gene was independently confirmed by qRT PCR.

### B1 B cells have high levels of glycolysis and oxidative phosphorylation

To confirm that enhanced expression of glycolysis genes had functional effects in B1a B cells, we measured glucose uptake and utilisation *ex vivo*. Transport of glucose through Glut1 is a key regulator of glycolytic rate, and has been shown to be critical in T-cell homeostasis (Macintyre et al., 2014). To quantify expression of this transporter we measured the binding of an eGFP-tagged HTLV-1 fusion protein which specifically binds to Glut1 (Manel et al., 2003). We found that B1a B cells have increased surface levels of the Glut1 transporter compared with Fo B2 and CD23^+^ peritoneal B2 cells, which do not express CD11b and are transiting the peritoneum, and splenic B1a B cells, which share the same microenvironment as Fo B2 B cells (Figure 2A). To exclude the possibility that the differences we observed were simply due to unequal cell size, we analysed eGFP fluorescence in peritoneal B1a and follicular B2 B cells gated to equalise flow cytometric forward scatter. Significant differences between the cell types remained (Supplementary Figure 1A). We next quantified glucose uptake using the fluorescent D-glucose analogue 2-NBDG. 2-NBDG was taken up more avidly in B1a B cells than Fo B2 cells (Figure 2B). Finally, we directly measured the basal extracellular acidification rate, which approximates to glycolytic flux, using Seahorse (Figure 2C). This confirmed high levels of aerobic glycolysis in B1 cells compared with Fo B2 cells. The second major source of ATP generation is by mitochondrial OXPHOS. We also noted elevated rates of basal oxygen consumption, reflective of active oxidative phosphorylation (OXPHOS) in B1 cells (Figure 2D).

**2.**
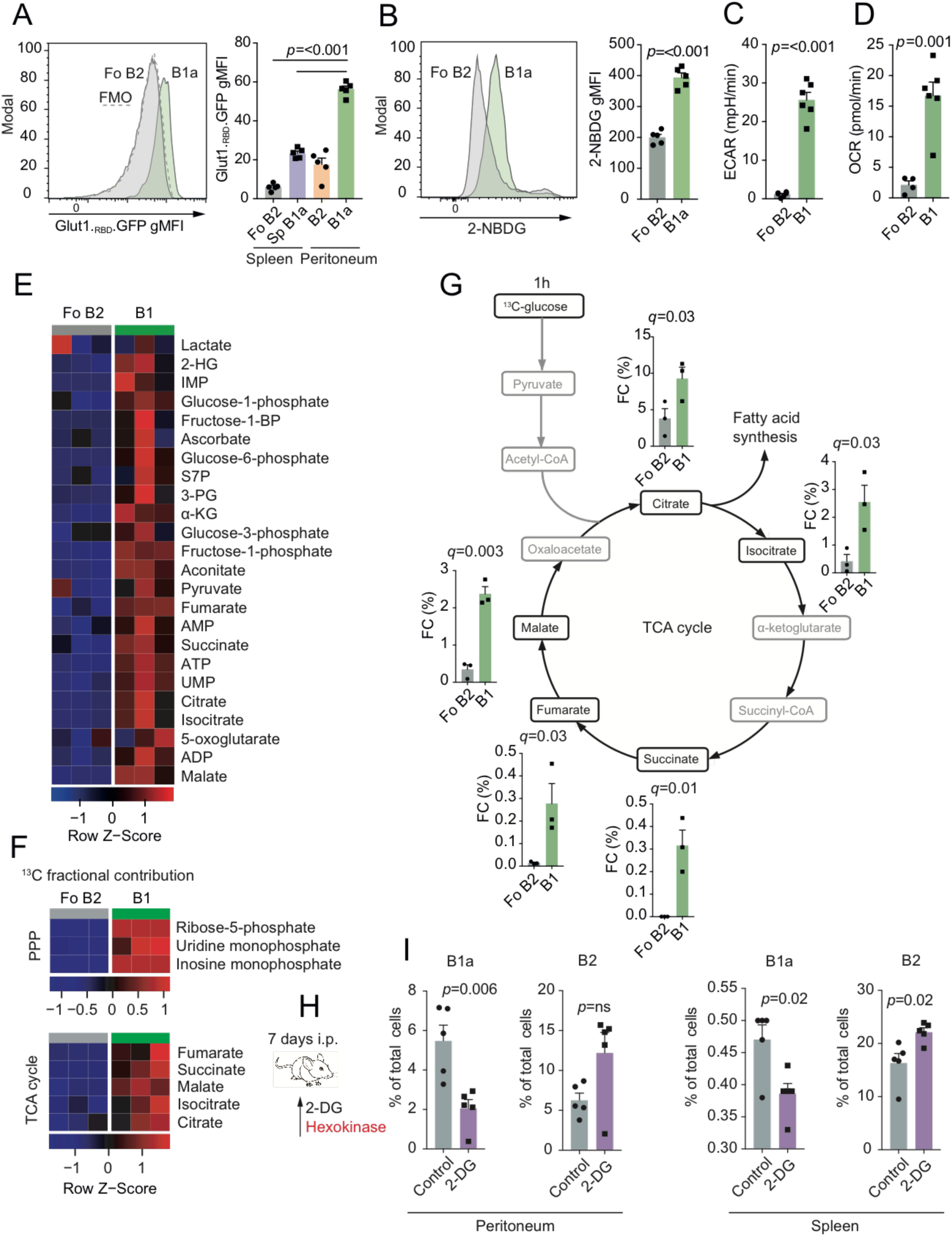
B1a B cells have active glycolysis and OXPHOS and low metabolic flexibility *in vivo*. **A**. Representative histogram of Glut1._RBD_.GFP staining in peritoneal B1a and Fo B2 B cells, and quantification of geometric mean fluorescence (gMFI) of Glut1._RBD_.GFP staining in B1a and B2 cells from the spleen and peritoneum. n=5 wild-type C57BL/6 individual mice per group. Data representative of two independent experiments. Mean ±SEM is depicted. One-way ANOVA with Dunnett correction for multiple testing used. **B**. Representative histogram of 2-NBDG uptake in peritoneal B1a and Fo B2 B cells Geometric mean fluorescence of 2-NBDG in splenic follicular B2 B cells and peritoneal B1a B cells. n=5 wild-type C57BL/6 individual mice per group. Data representative of two independent experiments. Mean ±SEM is depicted. Unpaired *t*-test used. **C**. Basal extracellular acidification rate (ECAR) of peritoneal total B1 B cells (B1a and B1b) compared with follicular B2 B cells. Data are n=5 technical replicates of cells sorted from pools of 10 wild-type C57BL/6 mice. Mean ±SEM is depicted. Unpaired *t*-test used. Data representative of two independent experiments. **D**. Basal oxygen consumption rate (OCR) of peritoneal total B1 B cells (B1a and B1b) compared with follicular B2 B cells. Data are n=5 technical replicates of cells sorted from pools of 10 wild-type C57BL/6 mice. Mean ±SEM is depicted. Unpaired *t*-test used. Data representative of two independent experiments. **E**. Heatmap of relative abundance of polar metabolites extracted from peritoneal total B1 (CD19^+^CD23^−^) and splenic follicular B2 B cells (CD19^+^CD23^+^) sorted by flow cytometry. Each data point represents cells sorted from pools of 5-7 6-week old wild-type C57BL/6 mice. Data representative of two independent experiments. Heatmap colouring is based on row Z-score. Abbreviations: 2-HG - 2-hydroxyglutarate; IMP - inosine monophosphate; S7P – sedoheptulose-7-phosphate; 3-PG – 3-phosphoglycerate; α-KG - α-ketoglutarate; UMP – uracil monophosphate. **F**. Heatmap of fractional contribution of ^13^C to metabolites in the PPP and the TCA cycle following incubation of cells described in (**E)** with [U-^13^C-glucose] for 1 hour. Data representative of two independent experiments. Heatmap colouring is based on row Z-score. **G**. Fractional contribution of ^13^C to metabolites in the TCA cycle following incubation of cells described in (**E)** with [U-^13^C-glucose] for 1 hour. Data representative of two independent experiments. Mean ±SEM is depicted. Unpaired *t*-test p-value is adjusted for multiple testing using FDR method (5% threshold). **H**. Experimental schematic. Wild type C57BL/6 mice were injected *i.p*. with 2-DG or PBS (control) each day for 7 days. The mechanistic target of 2-DG is illustrated in red.

Having demonstrated higher glucose uptake in B1a B cells in association with increased OXPHOS, we next traced carbon distribution from glucose using uniformly labelled ^13^C-glucose (i.e. all carbons are of the ^13^C isotope), followed by ion chromatography-mass spectrometry metabolomics. We incubated *ex vivo* peritoneal total B1 (CD19^+^CD23^−^) and Fo B2 B cells in ^13^C-glucose-supplemented media for one hour without stimulation, to maximise fidelity to *in vivo* metabolism.

We found that B1 B cells had elevated total levels of glycolytic, pentose phosphate pathway (PPP), and tricarboxylic acid (TCA) cycle intermediates compared to Fo B2 B cells (Figure 2E). The PPP runs parallel to glycolysis, and has anabolic functions, including the provision of ribose-5-phosphate for nucleic acid synthesis, and nicotinamide adenine dinucleotide phosphate (NADPH) for reductive reactions such as fatty acid synthesis. There was substantially more glucose-derived ^13^C contributing to TCA cycle and PPP metabolites in B1 B cells than Fo B2 B cells (Figure 2F).

Having demonstrated high levels of glycolysis and TCA cycle activity, we next determined the importance of glycolysis for B1 B cell homoeostasis *in vivo*. To do so, we treated mice with the compound-2-deoxyglucose (2-DG), which competitively inhibits glycolysis (Figure 2H-I). We found that 7 days of 2-DG treatment selectively depleted B1a B cells in both the peritoneum and spleen. 2-DG induced primary necrosis of B cells in the peritoneum, as determined by increased late necrotic cell numbers, but reduction in caspase-3 activation (Supplementary Figure 1B-C). Apoptosis was however activated in splenic B1a B cells (Supplementary Figure 1B). There was no effect of 2-DG on levels of the cell proliferation marker Ki67 in peritoneal B1a B cells, but a decrease was seen in splenic Fo B2 B cells (Supplementary Figure 1D). The lack of Ki67 upregulation in peritoneal B1a B cells suggested a failure to undergo homeostatic proliferation in the face of niche depletion.

To understand whether stimulation might influence relative pathway dependence, and to confirm our *in vivo* observations, we cultured B1 or Fo B2 B cells for 24 hours in the presence or absence of the TLR9 agonist ODN1826, with or without 2-DG. We found that 2-DG increased cell death two-fold in B1 B cells compared with Fo B2 B cells (Supplementary Figure 1E). IgM production was severely reduced in both cell types (Supplementary Figure 1F).

This result demonstrates that B1a B cells are dependent on glycolysis *in vivo*. Our finding that the proto-oncogene c-Myc is highly expressed in B1a B cells (Figure 1D), also recently reported by Hayakawa et al. (Hayakawa et al., 2016), suggests a driving mechanism for this pattern of energy metabolism, as c-Myc is a positive regulator of both glycolysis and oxidative phosphorylation (Caro-Maldonado et al., 2014). Notably, mice overexpressing c-Myc in B cells have increased B1 B cell numbers (Khuda et al., 2008).

### B1 B cells take up exogenous lipids and undergo cell death in response to inhibition of fatty acid synthesis

The finding of high levels of glucose uptake and entry into the TCA cycle and PPP pathway was suggestive of fatty acid synthesis, and given the residence of B1a B cells in the highly lipid rich peritoneal microenvironment, we next considered whether B1a B cells might also acquire exogenous lipids.

We injected mice intravenously with a fluorescently-labelled form of the long-chain fatty acid palmitate (BODIPY FL C_16_), and analysed them after one hour (Figure 3A-B). We found that peritoneal B1a B cells have increased fatty acid uptake compared to peritoneal CD23^+^ B2 cells, a finding preserved in the spleen, suggesting that a high capacity to take up free fatty acids is a persistent attribute of B1a B cells regardless of their local environment.

**3.**
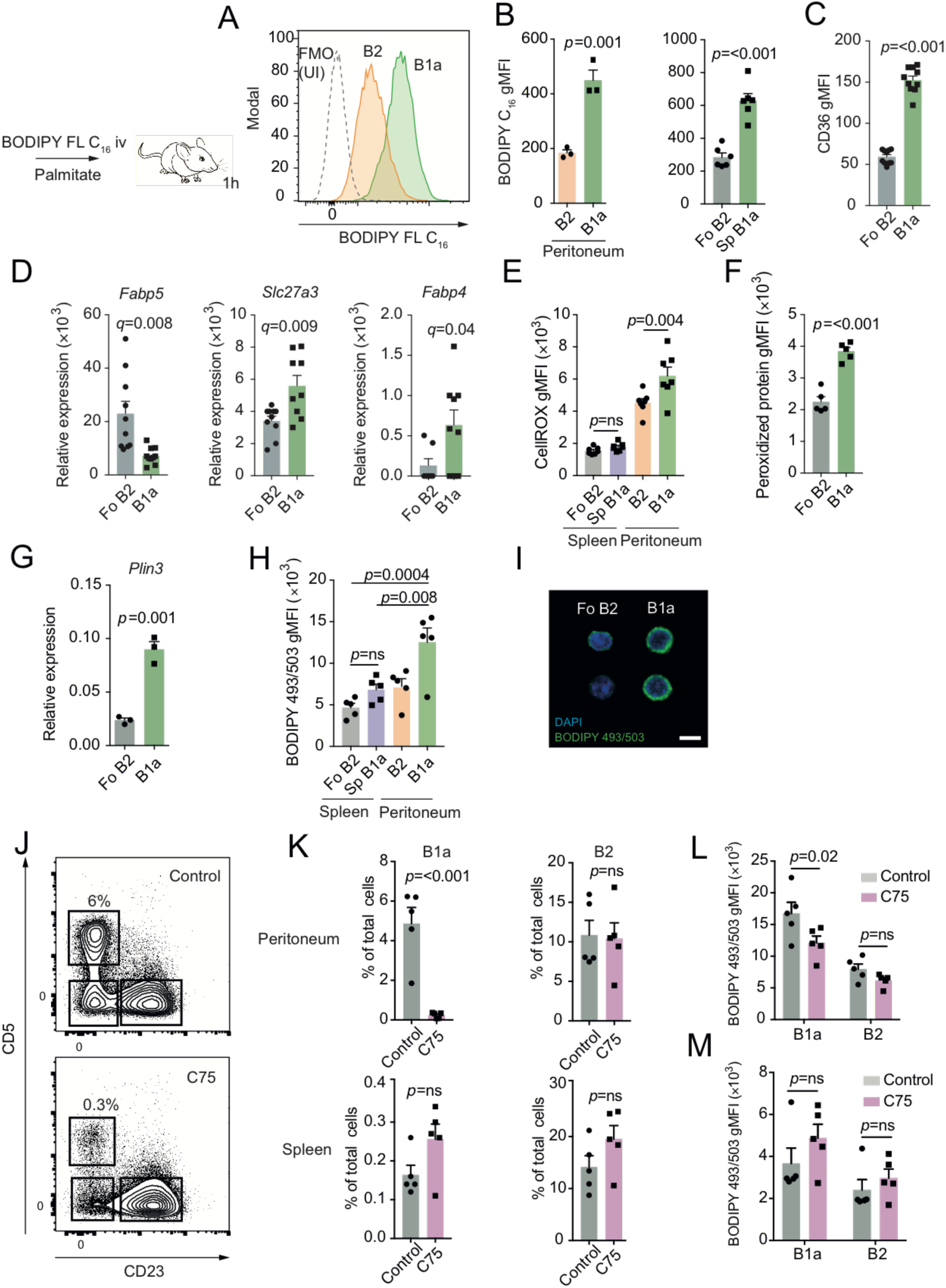
B1 B cells take up exogenous lipids and undergo cell death in response to inhibition of fatty acid synthesis. **A**. Experimental schematic and example histograms. 6-week-old C57BL/6 mice were injected intravenously with BODIPY FL C_16_ then analysed after 1 hour. Example distributions of fluorescence from peritoneal B1a and B2 B cells analysed by flow cytometry, compared with an uninjected control. **B**. Geometric mean fluorescence intensity of BODIPY FL C_16_ from peritoneal B cells following injection. n=3 biological replicates representative of or pooled from two independent experiments. Mean ±SEM is depicted. Unpaired *t*-test used. **C**. Geometric mean fluorescence intensity of CD36 on peritoneal B1a and splenic follicular B2 B cells from 6-week-old C57BL/6 mice. Each point represents a single mouse, and data are pooled from two independent experiments. Mean ±SEM is depicted. Unpaired *t*-test used. **D**. Peritoneal B1a and splenic follicular B2 B cells were sorted by flow cytometry from 6-week-old C57BL/6 mice and qRT-PCR was performed for the indicated genes, relative to *B2m*. Each point represents a single mouse, and data are pooled from two independent experiments. Mean ±SEM is depicted. Unpaired *t*-test *p*-value is adjusted for multiple testing using FDR method (5% threshold). **E**. Geometric mean fluorescence intensity of CellROX from splenic and peritoneal B cells of 6-week-old C57BL/6 mice. n=5 biological replicates. Mean ±SEM is depicted. One-way ANOVA with Dunnett correction for multiple comparison used. n=5 biological replicates. Representative of two independent experiments. **F**. Geometric mean fluorescence intensity of Alexa Fluor 488-ClickIT lipid peroxidation assay for peritoneal B1a and splenic Fo B2 B cells of 6-week-old C57BL/6 mice. n=5 biological replicates. Mean ±SEM is depicted. Unpaired *t*-test used. Representative of two independent experiments. **G**. Expression of *Plin3* relative to *B2m* in flow sorted peritoneal B1a and splenic follicular B2 B cells from 6-week-old C57BL/6 mice. n=3 biological replicates. Mean ±SEM is depicted. Unpaired *t*-test used. Representative of two independent experiments. **H**. Geometric mean fluorescence intensity of BODIPY 493/503 in splenic follicular B2 B cells, and peritoneal B1a and B2 B cells of 6-week-old C57BL/6 mice. n=5 biological replicates, pooled from two independent experiments. Mean ±SEM is depicted. One-way ANOVA with Dunnett correction for multiple comparison used. **I**. Representative microscope images of BODIPY 493/503 staining in flow sorted peritoneal B1a and splenic follicular B2 B cells (x60 magnification) 6-week-old C57BL/6 mice. Scale bar is 20μm. Representative of two independent experiments. **J-M**. 6-week-old C57BL/6 mice received *i.p*. injections of C75 (15mg.kg^−1^) every 2 days for 3 doses, before sacrifice. Shown are representative flow cytometry plots of peritoneal CD19^+^ B cells from C75 injected and control mice (H). Shown is the percentage of the indicated population of total cells in the analysed compartment (I) or the geometric mean fluorescence intensity of BODIPY 493/503 measured by flow cytometry (J-K). Each point represents one mouse. Mean ± SEM is depicted. Unpaired *t*-test used in (I), two-way ANOVA with Sidak correction for multiple testing used in (J) and (K). Representative of 3 independent experiments.

To analyse how these lipids might be taken up, we next quantified surface expression of the fatty acid transporter protein CD36 by flow cytometry. We found approximately three-fold higher levels of CD36 on peritoneal B1a B cells compared with Fo B2 B cells (Figure 3C). We also examined expression of other lipid transporter genes by qPCR, and found higher levels of *Slc27a3* and *Fabp4* transcription in B1a B cells, but decreased *Fabp5* expression compared to Fo B2 B cells, suggesting subset-specific lipid transporter use (Figure 3D).

The presence of high concentrations of intracellular free fatty acids can lead to oxidative stress caused by lipid peroxidation (Hauck and Bernlohr, 2016). In keeping with this, we found elevated levels of cellular reactive oxygen species (ROS) and lipid peroxidation-derived protein modifications in B cells from the peritoneum compared with the spleen, and a relative increase in these parameters in the peritoneal B1a compared with B2 populations (Figure 3E-F), suggesting that the peritoneal microenvironment inherently induces oxidative stress in association with lipid flux. In order to mitigate lipotoxicity, triglycerides are synthesised from fatty-acyl CoA, which are then stored in lipid droplets (Wilfling et al., 2014). The finding that expression of the lipid droplet-associated gene *Perilipin-3* is increased in B1a B cells suggests that they may store excess lipid in droplet form (Figure 3G). To explore this, we measured neutral lipid storage by staining with the BODIPY493/503 tracer molecule. By flow cytometry and microscopy, we found significantly more lipid droplets in B1a B cells compared with Fo B2 cells (Figure 3H-I).

To directly assess the importance of fatty acid synthesis to the maintenance of B1a B cells, we treated mice with 3 doses of the fatty acid synthase (FAS) inhibitor C75 on alternate days. C75 treatment led to a dramatic depletion of peritoneal B1a B cells, with no significant effect on peritoneal B2 B cells, or on either subset in the spleen (Figure 3J-K). In keeping with its mode of action, neutral lipid stores were depleted in peritoneal B1a B cells but not peritoneal B2 B cells, or in splenic populations (Figure 3L-M). The lack of impact on B2 B cells is in accordance with previous work *in vitro*, which showed no effect of C75 on splenic B cells (Bhatt et al., 2012). To analyse how C75 might lead to B1 B cell depletion, we analysed caspase-3 activation and levels of the proliferation marker Ki67 by intracellular flow cytometry following the 3-dose regime. We found that Ki67 levels were lower in the splenic B1a B cells of C75 treated mice, but no differences were observed in peritoneal B cell subsets or in caspase-3 activation (Supplementary Figure 2A-B). This suggests that in the peritoneum, B1 cell loss occurs rapidly, and that the expected increase in Ki67 as B1a B cells expand to fill the deficient niche does not occur.

These results together indicate that B1a B cells both acquire exogenous lipids, and also require fatty acids synthesised *de novo*, which is likely to be from citrate produced as a product of glycolysis, through the enzyme ATP-citrate lyase.

### Autophagy is required for B1 B cell survival and self-renewal

Autophagy is a highly-conserved process for the degradation of cellular macromolecules and organelles by the lysosome (Levine et al., 2011). This pathway is therefore capable of supplying critical substrates to fuel metabolism (Kaur and Debnath, 2015; Riffelmacher et al., 2017).

Autophagy has been shown to be required for the control of cellular lipid dynamics, with roles both in degradation and lipid droplet formation (Liu and Czaja, 2012; Singh et al., 2009; Rambold et al., 2015). Active autophagy is a key attribute maintaining the metabolic state required for self-renewal (Mortensen et al., 2011; Pan et al., 2013; Guan et al., 2013; García-Prat et al., 2016). In the resting state, deletion of autophagy genes has been reported to selectively lead to loss of B1 B cells (Miller et al., 2008). We reasoned that autophagy may therefore be critical in maintaining normal B1 B cell metabolic homeostasis by the provision of metabolites such as free fatty acids, which would also therefore affect their self-renewal capacity.

We first measured autophagic flux by washing out non-autophagosomal LC3-I with saponin and then using immunofluorescence to detect LC3-II by flow cytometry (Figure 4A) (Eng et al., 2010). This assay was performed with and without bafilomycin A_1_, a lysosomal V-ATPase inhibitor which prevents autophagosome-lysosome fusion and therefore reveals autophagic flux. We found considerably increased autophagy in peritoneal B1a B cells compared with Fo B2 B cells. We next conditionally deleted *Atg7* in B cells using the *Mb1*-Cre system (hereafter referred to as B-*Atg7*^−/−^) (Hobeika et al., 2006). In keeping with other reports of B-cell autophagy deletion at later points in B-cell development, we found an essentially normal peripheral B cell compartment, but markedly reduced numbers of peritoneal B1 B cells (Figure 4B and Supplementary 3A) (Miller et al., 2008; Chen et al., 2014; Arnold et al., 2015). Also reduced were splenic B1a B cells (Figure 4C) and interestingly, peritoneal B2 B cells (Figure 4D).

**4.**
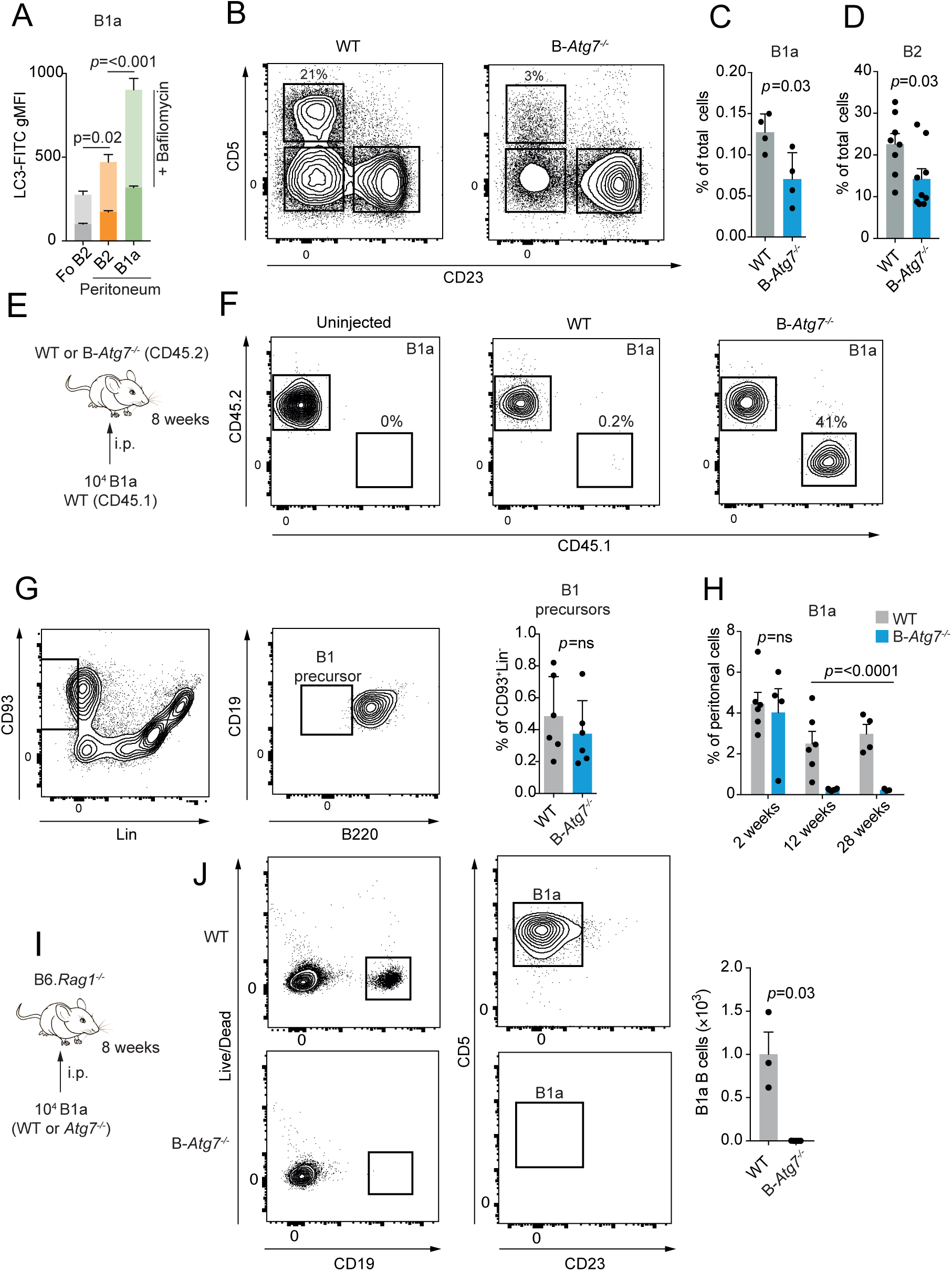
Autophagy is required for B1a survival and self-renewal. **A**. Geometric mean fluorescence intensity of LC3-FITC in splenic follicular B2 and peritoneal B1a and B2 B cells following wash-out of LC3-I. Overlay bars denote gMFI following incubation with 10nM bafilomycin A_1_ for 30 minutes. n=5 biological replicates pooled from two independent experiments. Mean±SEM is depicted. One-way ANOVA with Dunnett correction for multiple testing calculated on post-bafilomycin values. **B**. Representative example of peritoneal CD19^+^ B cell compartment of control and B-*Atg7*^−/−^ mice. **C**. Splenic B1a B cells as a percentage of total splenic cells in control and *B-Atg7*^−/−^ mice. Each point represents one mouse. Mean ±SEM is depicted. Unpaired *t*-test used. Representative of >4 independent experiments. **D**. Peritoneal B2 B cells as a percentage of total peritoneal cells in control and *B-Atg7*^−/−^ mice. Each point represents one mouse. Mean ±SEM is depicted. Unpaired *t*-test used. Representative of >4 independent experiments. **E**. Experimental schematic. 10^4^ sorted peritoneal B1a B cells from CD45.1 wild-type mice were transferred by *i.p*. injection to either control or *B-Atg7*^−/−^ mice (both CD45.2). Representative of 2 independent experiments. **F**. Representative flow cytometric plot of peritoneal cells from **E**. **G**. Gating strategy for bone marrow B1-precursors. Cells were defined as viable, CD93^+^Lin^−^CD19^+^B220^−^. B1 precursor cells as a percentage of total bone marrow cells in control and *B-Atg7*^−/−^ mice are shown. Each point represents one mouse. Mean ±SEM is depicted. Representative of 2 independent experiments. **H**. Peritoneal B1a B cells as a percentage of total peritoneal cells in control and *B-Atg7*^−/−^ mice at 2 weeks, 12 weeks, and 28-weeks of age. Each point represents one mouse. Data pooled from two independent experiments. Mean ±SEM is depicted. Two-way ANOVA with Sidak correction for multiple comparison used. **I**. Experimental schematic. 10^4^ sorted peritoneal B1a B cells from control and B-*Atg7*^−/−^ mice were adoptively transferred by *i.p*. injection to B6.*Rag1*^−/−^ hosts, and analysed after 8 weeks. **J**. Representative flow cytometric plot of peritoneal cells from **I**. Each point represents one mouse. Mean ±SEM is depicted. Unpaired *t*-test used. Representative of 2 independent experiments.

B1a cells once stimulated leave the peritoneum via the omentum and traffic to the spleen and bone marrow to produce immunoglobulin (Ansel et al., 2002; Yang et al., 2007). Since autophagy is required for plasma cell formation and maintenance (Pengo et al., 2013; Clarke et al., 2014), it was possible that B1a B cells might rapidly exit the peritoneum in an attempt to regulate natural IgM levels in a *B-Atg7*^−/−^ environment. To determine if this was the case, we next adoptively transferred wild-type congenically marked CD45.1 B1a B cells into the peritonea of *B-Atg7*^−/−^ mice (which express the CD45.2 antigen) (Figure 4E-F). We found that wild-type CD45.1 B1a B cells remained in the peritoneum and effectively self-renewed following transfer, excluding the possibility that autophagy deficient B1a B cells were simply dispersing. Next, to determine if the B1 B cell defect was due to failure of normal development, we examined the numbers of CD93^+^Lin^−^CD19^+^B220^+^ B1 precursor cells in B-*Atg7*^−/−^ mice (Figure 4G), which are generated throughout life at a low level from the bone marrow (Esplin et al., 2009). We found no difference in these progenitors compared with control mice, suggesting that their maintenance does not require autophagy. To examine the kinetics of B1 B cell loss, we analysed mice at 2 weeks, 12 weeks, and 28 weeks of age (Figure 4H). We found no difference in B1a numbers at 2 weeks, but by 12 weeks they had drastically decreased, and remained low at 28 weeks, indicating that B1 B cell fetal and neonatal differentiation is normal, but self-renewal in adult mice is affected by loss of autophagy. To confirm that autophagy is required for B1a B cell self-renewal, we adoptively transferred 10^4^ sorted autophagy-deficient or control cells into the peritonea of B6. *Rag1*^−/−^ hosts (Figure 4I-J). After 8 weeks, *B-Atg7*^−/−^ B1a B cells were undetectable, in contrast to control cells which were now abundant. These results therefore demonstrate a cell intrinsic defect in self-renewal of B1a B cells when autophagy is absent.

### Autophagy maintains B1 B cell metabolic homeostasis

To investigate the impact of autophagy deficiency on peritoneal B1a and splenic Fo B2 B cell metabolism, we repeated multiplex qRT-PCR with Fluidigm Biomark (Figure 5A). There was a generalised reduction in the expression of metabolism associated genes in B1a B cells, which was much less pronounced in Fo B2 B cells. Notably downregulated was Acaca, which encodes the key regulator of fatty acid synthesis, acetyl-CoA carboxylase 1. Post-hoc analysis of other lipid metabolic genes also revealed downregulation of *Acacb*, *Acsl1*, *Plin3*, *and Srebf2* (Supplementary Figure 3B). Expression of these genes was not affected by autophagy deficiency in Fo B2 B cells, and no other genes were significantly differentially expressed following adjustment for multiple testing. *Atg7* was effectively deleted in B-*Atg7*^−/−^ B1a B cells, excluding the possibility that the residual population was due to failure of Cre recombinase activity (Supplementary Figure 3C).

**5.**
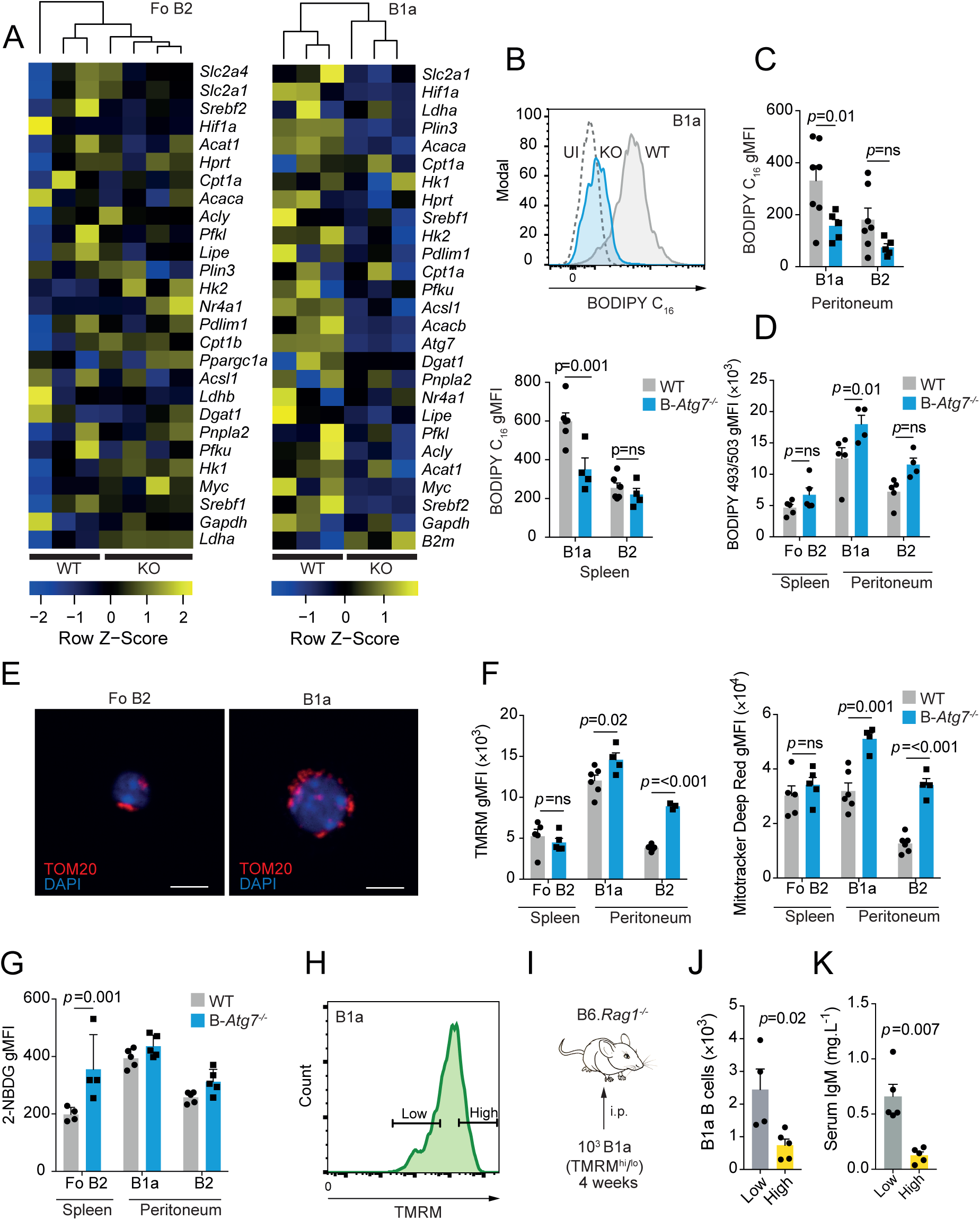
**Loss of autophagy leads to disrupted B1a B cell metabolic homeostasis** (Legend continues on next page) **A**. Heatmap of Fluidigm Biomark qRT-PCR gene expression data for peritoneal B1a B cells and follicular splenic B2 B cells (CD19^+^CD23^+^) from control and B-*Atg7*^−/−^ mice. Data are relative to β-actin, and colouring is based on row Z-score. Each data point is the mean of three technical replicates. Hierarchical clustering is unsupervised. **B**. Example distributions of fluorescence of BODIPY FL C_16_ from peritoneal B1a B cells from control and B-*Atg7*^−/−^ mice, measured by flow cytometry, compared with an uninjected wild-type control. **C**. Geometric mean fluorescence of BODIPY FL C_16_ in B1a and B2 B cells from peritoneum and spleen, in control and B-*Atg7*^−/−^ mice, measured by flow cytometry. Each point represents one mouse. Mean ±SEM is depicted. Data pooled from two independent experiments. Two-way ANOVA with Sidak correction for multiple testing used. **D**. Geometric mean fluorescence of BODIPY 493/503 in B1a and B2 B cells from peritoneum and spleen, in control and B-*Atg7*^−/−^ mice, measured by flow cytometry. Each point represents one mouse. Mean ±SEM is depicted. Data pooled from two independent experiments. Two-way ANOVA with Sidak correction for multiple testing used. **E**. Representative immunofluorescent confocal images of the mitochondria of sorted wild type follicular B2 and peritoneal B1a B cells. Cells are stained for TOM20 (red) and nuclei are visualised with DAPI (blue). 60× magnification, scale bar is 5μm. **F**. Geometric mean fluorescence of TMRM and Mitotracker Deep Red in B cells from the spleen and peritoneum, in control and B-*Atg7*^−/−^ mice, measured by flow cytometry. Each point represents one mouse. Mean ±SEM is depicted. Data representative of two independent experiments. Two-way ANOVA with Sidak correction for multiple testing used. **G**. Geometric mean fluorescence of 2-NBDG in B1a and B2 B cells from peritoneum and spleen, in control and B-*Atg7*^−/−^ mice, measured by flow cytometry. Each point represents one mouse. Mean ±SEM is depicted. Data pooled from two independent experiments. Two-way ANOVA with Sidak correction for multiple testing used. **H**. Flow cytometry gating definition of high and low levels of TMRM fluorescence. The highest and lowest quartiles of the distribution were used to define TMRM^hi^ and TMRM^lo^ populations of B1a B cells. **I**. Experimental schematic. 10^3^ B1a B cells from the lowest and highest quartiles of TMRM fluorescence were sorted and adoptively transferred into the peritonea of B6.*Rag7*^−/−^ hosts. These mice were then analysed after 1 month. **J**. Quantification of peritoneal B1a B cells from **I**. Each point represents one mouse. Mean ±SEM is depicted. Unpaired *t*-test used. Data representative of two independent experiments. **K**. Quantification of serum IgM levels from **I**. Each point represents one mouse. Mean ±SEM is depicted. Unpaired *t*-test used. Data representative of two independent experiments.

The downregulation of genes involved in fatty acid synthesis by autophagy-deficient B1a B cells suggested that lipid uptake and storage might be defective. To understand how autophagy deficiency affected lipid homeostasis, we first intravenously injected control and B-*Atg7*^−/−^ mice with BODIPY FL C_16_ (Figure 5B-C). We found that exogenous fatty acid uptake was reduced in B1a B cells from the peritoneum or spleen in B-*Atg7*^−/−^ mice, but Fo B2 cells were unaffected. However, total neutral lipid stores were in fact increased in autophagy-deficient peritoneal B1a B cells, but Fo B2 cells were again unaffected (Figure 5D). These results implied that loss of autophagy inhibited fatty acid uptake in the context of accumulation of intracellular lipids, and based on transcriptional data, was associated with a decrease in lipid synthesis. The increase in lipid content of B-*Atg7*^−/−^ B1a B cells can be explained by loss of lipophagy, a process by which lipid droplets are degraded by autophagy to release free fatty acids (Singh et al., 2009). The transcriptional switch away from lipid synthesis may reflect an attempt to reduce potentially toxic accumulation of intracellular lipids.

To assess to what extent mitochondrial OXPHOS might be affected by loss of autophagy, we first visualised mitochondria from Fo B2 and peritoneal B1a B cells by confocal immunofluorescent microscopy (Figure 5E). B1a B cells had a more extensive mitochondrial network than Fo B2 B cells, whose mitochondria were punctate. The fused mitochondrial pattern seen in B1a B cells is reported in cells with active OXPHOS, but also in those undergoing stress, which could include the lipid toxicity previously noted (Figure 3F) (Youle and van der Bliek, 2012; Buck et al., 2016). We next quantified mitochondrial membrane potential (MMP) using tetramethylrhodamine (TMRM), a fluorescent dye sequestered by active mitochondria, and which is reflective of electron transfer chain activity. We found that in wild-type mice, TMRM fluorescence was significantly increased in B1a B cells compared with either Fo B2 or peritoneal B2 B cells (Figure 5F). However, total mitochondrial mass, measured using Mitotracker Deep Red, was similar in Fo B2 B cells and peritoneal B1a B cells, but much lower in peritoneal B2 B cells. These results suggested therefore, that in the peritoneal microenvironment, mitochondrial activity and therefore OXPHOS is increased, which may reflect fatty acid availability as a fuel source. When we compared Fo B2 B cells of B-*Atg7*^−/−^mice with wild-type, we found no difference in mitochondrial mass or TMRM intensity (Figure 5F). Sparing of Fo B2 B cell mitochondrial function may therefore reflect either their intrinsic metabolic program, or the difference in microenvironment.

We next assessed whether there was a compensatory increase in glycolysis in autophagy-deficient B cells (Figure 5G), as has been demonstrated in other cell types (Kabat et al., 2016; Watson et al., 2015; Puleston et al., 2014). Peritoneal B1a B cells failed to increase their uptake of 2-NBDG. However, *Atg7*^−/−^ Fo B2 B cells did significantly upregulate their glucose uptake. B1a B cells therefore have limited metabolic plasticity when autophagy is lost, unlike Fo B2 B cells, which are able to compensate by upregulation of glycolytic activity. This lack of flexibility is in keeping with in vivo inhibitor data (Figure 2I). Direct measurement of metabolite levels, or assessment of extracellular and metabolic flux was not possible in *Atg7*^−/−^ B1a B cells, due to their severely reduced numbers.

Recent work has shown that MMP is an important determinant of the capacity of a number of cell types, including HSCs and CD8^+^ T cells, to self-renew (Sukumar et al., 2016; Vannini et al., 2016). CD8^+^ T cells with low MMP have increased free fatty acids and lower rates of OXPHOS, and an enhanced capacity to self-renew following adoptive transfer (Sukumar et al., 2016). Moreover, autophagy was found to be more active in HSCs which had undergone mitochondrial depolarisation (Vannini et al., 2016). After finding that loss of autophagy in peritoneal B1a B cells resulted in an increase in MMP, we hypothesised that an increase in hyperpolarised mitochondria might mechanistically contribute to the self-renewal defect seen when autophagy is deleted. We therefore sorted wild-type peritoneal B1a B cells with high and low levels of MMP and adoptively transferred these subsets into B6.*Rag1*^−/−^ mice (Figure 5H-J). There were significantly fewer B1a B cells detected in the peritonea of mice which had received cells with high MMP compared with those that had received the low MMP subset, and serum natural IgM levels were also accordingly lower (Figure 5K). These data therefore suggest that accumulation of active mitochondria is associated with the failure of self-renewal seen in B cells that lack autophagy.

We have demonstrated that B1a B cells engage a specific, autophagy-dependent, metabolic program to survive and self-renew in the peritoneal microenvironment, characterised by lipid uptake and predominant fatty acid synthesis, but also high levels of glycolysis, PPP, and TCA activity. The evolution of this pattern of metabolism is in keeping with the ability of B1 B cells to rapidly respond to infection, compared with B2 B cells. Interestingly, this activated state in also seen in cells which have undergone malignant transformation (Bhatt et al., 2012). Many forms of leukaemia are CD5^+^, and early-generated B1-like B cells have been demonstrated to be a source of chronic lymphocytic leukaemia (CLL) (Hayakawa et al., 2016). CLL has been found to have many metabolic features in common with our observations in B1 B cells, including high rates of lipid storage, glycolysis, and oxidative phosphorylation (Jitschin et al., 2014; Doughty, 2006; Tili et al., 2012; Rozovski et al., 2016).

Understanding whether the physiological metabolic phenotype we have described in B1 B cells contributes to their malignant potential remains an area for further study, as does the potential for inhibition of fatty acid synthesis or autophagy as a novel therapeutic target.

### Author contributions

A.C. and A.K.S. conceived the study and wrote the paper. A.C., T.R., and D.B. designed and performed experiments and analysed data. R.J.C. designed experiments and analysed data.

## Acknowledgements and funding

We thank the staff of the University of Oxford Biomedical Services Unit for animal care, Jonathan Webber and Craig Waugh for assistance with FACS sorting, Mariolina Salio for assistance with Seahorse, Zhanru Yu for technical assistance, and Fiona Powrie for providing B6.*Rag7*^−/−^ mice. A.C. is funded by a Wellcome Trust Clinical Postdoctoral Fellowship (104549/Z/14/Z) and A.K.S. is funded by a Wellcome Trust Investigator Award (103830/Z/14/Z). The authors have no competing financial interests.

## Methods

### Mice

*Atg7* was conditionally deleted in B cells by the generation of *Mb1*-cre × *Atg7^F/F^* mice(Hobeika et al., 2006). Littermate controls were used in comparative experiments. CD45.1 B6.SJL and C57BL/6 mice were purchased from Harlan, and acclimatised before use. B6.*Rag1*^−/−^ were supplied by Fiona Powrie (University of Oxford, UK) and bred in the local animal facility. All mice were housed in a specific pathogen free environment and used from between 6-12 weeks of age unless otherwise indicated. Male and female mice were used equally. Animal experiments were approved by the local ethical review committee and performed under UK Project License PPL 30/3388.

### Cell isolation and flow cytometry

Peritoneal cells were extracted by lavage. Splenic cells were prepared by straining through a 70μm mesh, then red blood cells lysed with Red Cell Lysis Buffer (Biolegend). For flow cytometry, dead cells were excluded using Fixable Viability Dye 780 (eBioscience), and Fc receptors blocked with anti-CD16/32 antibody (93, Biolegend). Fluorochrome conjugated anti-CD19 (6D5), anti-CD5 (53-7.3), anti-CD23 (B3B4), anti-IgM (RMM-1), anti-B220 (RA3-6B2), anti-CD45.1 (A20), and anti-CD45.2 (104), anti-CD93 (AA4.1, eBioscience), anti-Lineage cocktail (eBioscience), anti-CD21 (7E9), and anti-CD36 (HM36) antibodies were used for surface staining (all from Biolegend unless otherwise specified). Flow cytometry was performed on LSR II or Fortessa X20 instruments (BD Biosciences). Flow sorting was performed on a FACSAria III (BD Biosciences). Intracellular staining for anti-active caspase-3 (BD Biosciences, C92-605) and Ki67 (Biolegend, 11F6) was performed following fixation and permeabilisation (FoxP3/Transcription Factor Buffer Set, eBioscience).

### Adoptive transfer

B1a B cells were sorted from the peritoneal lavage of donor mice by flow cytometry using the gating strategy presented in Figure 1A. Cells were transferred by intraperitoneal injection.

### *In vivo* inhibitor treatment

Mice were treated with daily intraperitoneal injections of 2-deoxyglucose (1g.kg^−1^, Sigma), or PBS for 7 days, then sacrificed. C75 (15mg.kg^−1^, Cayman Chemical) was given three times per week, for one week. Doses and regimes were selected with reference to the literature(Wilhelm et al., 2016; Loftus et al., 2000).

### *In vitro* inhibitor treatment

Cells were cultured in RPMI 1640 medium supplemented with 10% FBS, 2 mM glutamine, and 50μM 2-mercaptoethanol (complete RPMI) and stimulated with 0.5μM ODN1826 (Miltenyi Biotec). 2-deoxyglucose was used at 2.5mM, selected with reference to the literature(Sukumar et al., 2013)

### BODIPY C_16_ uptake *in vivo*

Mice were injected intravenously with 50μg BODIPY FL C_16_ (Life Technologies) in 100µl PBS, then sacrificed after 1 hour.

### Neutral lipid staining

Cells were stained with BODIPY 493/503 (Life Technologies) at a concentration of 1μg.ml^−1^ in complete RPMI for 30 minutes at 37°C, then washed in PBS three times before analysis by flow cytometry or microscopy.

### Measurement of autophagy

Cells were incubated in complete RPMI for 30 minutes in the presence of 10nM bafilomycin A_1_ (Sigma), and then LC3 intracellular staining performed after permeabilization, using the FlowCellect LC3-antibody based assay kit (EMD Millipore) in accordance with manufacturer’s instructions.

### Confocal microscopy

Cells were sorted by flow cytometry and then fixed with Fixation Buffer (Biolegend). They were then permeabilised with 0.1% Triton X100 and blocked with Starting Block (Thermo). Primary antibody staining was with rabbit anti-TOM20 (Santa Cruz), and secondary Alexa Fluor 488 goat anti-rabbit antibody was used for visualisation. Cells were imaged with a Nikon TE2000 microscope. Image deconvolution was performed with ImageJ (NIH).

### Cellular reactive oxygen species and lipid peroxidation measurement

Cells were stained using CellROX Green (Life Technologies) at a concentration of 5μM at 37°C in complete RPMI for 30 minutes, then washed in PBS three times. For determination of lipid peroxidation, the Click-IT Lipid Peroxidation Kit AF488 (Life Technologies) was used as per manufacturer’s instructions.

### Mitochondrial dye assays

For MMP, cells were incubated with TMRM (Life Technologies) at a concentration of 25nM in complete RPMI for 30 minutes at 37°C, then washed in PBS three times. For mitochondrial mass, cells were incubated with Mitotracker Deep Red (Life Technologies) at a concentration of 100nM in complete RPMI for 30 minutes at 37°C, then washed in PBS three times.

### Glucose uptake

To measure surface Glut1 expression, cells were incubated with 2.5μL per test of Glut1._RBD_.GFP (Metafora Biosystems, France) for 20 minutes at 37ºC in complete media containing 0.09% sodium azide (Sigma), then analysed by flow cytometry. To measure glucose uptake, cells were incubated with 10μM 2-NBDG (Sigma) for 30 minutes at 37°C in complete media, then analysed by flow cytometry.

### Fluidigm Biomark

Peritoneal B1a and follicular B2 B cells were flow-sorted (200 cells/population) into OneStep lysis buffer (Invitrogen). RNA was reverse transcribed and cDNA was pre-amplified using the CellsDirect OneStep q-RT kit (Invitrogen). The selected metabolic genes (Methods Table 1) were amplified and analysed for expression using a dynamic 48×48 array (Biomark Fluidigm) as previously described by Tehranchi et al. (Tehranchi et al., 2010). Data were analyzed using the 2^−ΔCt^ method, and all results were normalized to β-actin, which was selected as the optimum housekeeping gene from the panel. Any genes not detected in both comparative populations were excluded from analysis. Heatmap generation, hierarchical clustering, and PCA were performed in R (3.2.3) using the package *gplots*. Gene expression differences were calculated by *t*-test, and *p*-values were adjusted using the method of Benjamini and Hochberg with the false discovery rate set to 5%.

### Mass spectrometry-based metabolomics analysis using ion chromatography

Three × 10^5^ peritoneal B1 B cells (CD19^+^CD23^−^) and splenic follicular B2 B cells (CD19^+^CD23^+^) were sorted by flow cytometry. Isolated cells were incubated in RPMI (glucose-free formulation) containing 10mM [U-^13^C]-glucose (Cambridge Isotope Laboratories, Canada), 2mM glutamine, and 10% dialysed fetal bovine serum (Thermofisher) at 37°C for 1 hour. Three × 10^5^ cells were washed in ice cold 150mM ammonium acetate (pH 7.3) and metabolites extracted in 80% methanol on dry ice before evaporation under vacuum. Dried metabolites were re-suspended in 50μl 50% ACN and 5μl were injected for chromatographic separation using the Thermo Scientific Ion Chromatography System (ICS) 5000 coupled to a Thermo Scientific Q Exactive run in negative polarity mode (as described in ((Nagaraj et al., 2017). The gradient ran from 5 mM to 95 mM KOH over 18 min with a flow rate of 350μl.min^−1^. The settings for the HESI-II source were: S-lens 50, Sheath Gas 18, Aux Gas 4, spray heater 320°C, and spray voltage −3.2 kV. Metabolites were identified based on accurate mass (± 3 ppm) and retention times of pure standards. Relative amounts, mass isotopologue distributions (MIDs) and fractional contributions of metabolites were quantified using TraceFinder 3.3. Heatmap generation and hierarchical clustering were performed in R (3.4.3) using the package *gplots*.

### Metabolic analysis with Seahorse extracellular flux analyser

Real-time extracellular acidification rate (ECAR) and oxygen consumption rate (OCR) were measured using a XFe 96 extracellular flux analyzer (Agilent). Cells were seeded at 2 × 10^5^ cells per well in Seahorse Base Medium supplemented with 10mM glucose, 2mM glutamine, and 1mM sodium pyruvate. Cells were rested for 1 hour at 37°C before analysis.

### ELISA

Total serum IgM was measured according to manufacturer’s instructions (Invitrogen)

### Quantitative real-time PCR

RNA was isolated using the RNeasy Micro Kit (Qiagen) and reverse transcribed using the High Capacity cDNA Reverse Transcription Kit (Thermo Scientific). Quantitative real-time PCR was carried out using Taqman Gene Expression Master Mix on a ViiA 7 instrument (Thermo Scientific). PCR probes used are listed in Supplementary Table 1.

### Statistics

Data was analyzed with Prism 7 (Graphpad). Two populations were compared by unpaired *t*-test. Three or more sample populations were compared by one-way ANOVA with multiple testing correction according to the method of Dunnett. Groups of populations were compared by two-way ANOVA with multiple testing correction using the method of Sidak. *P*-values were considered significant if <0.05.

## Supplementary figure

**Supplementary Figure 1.**
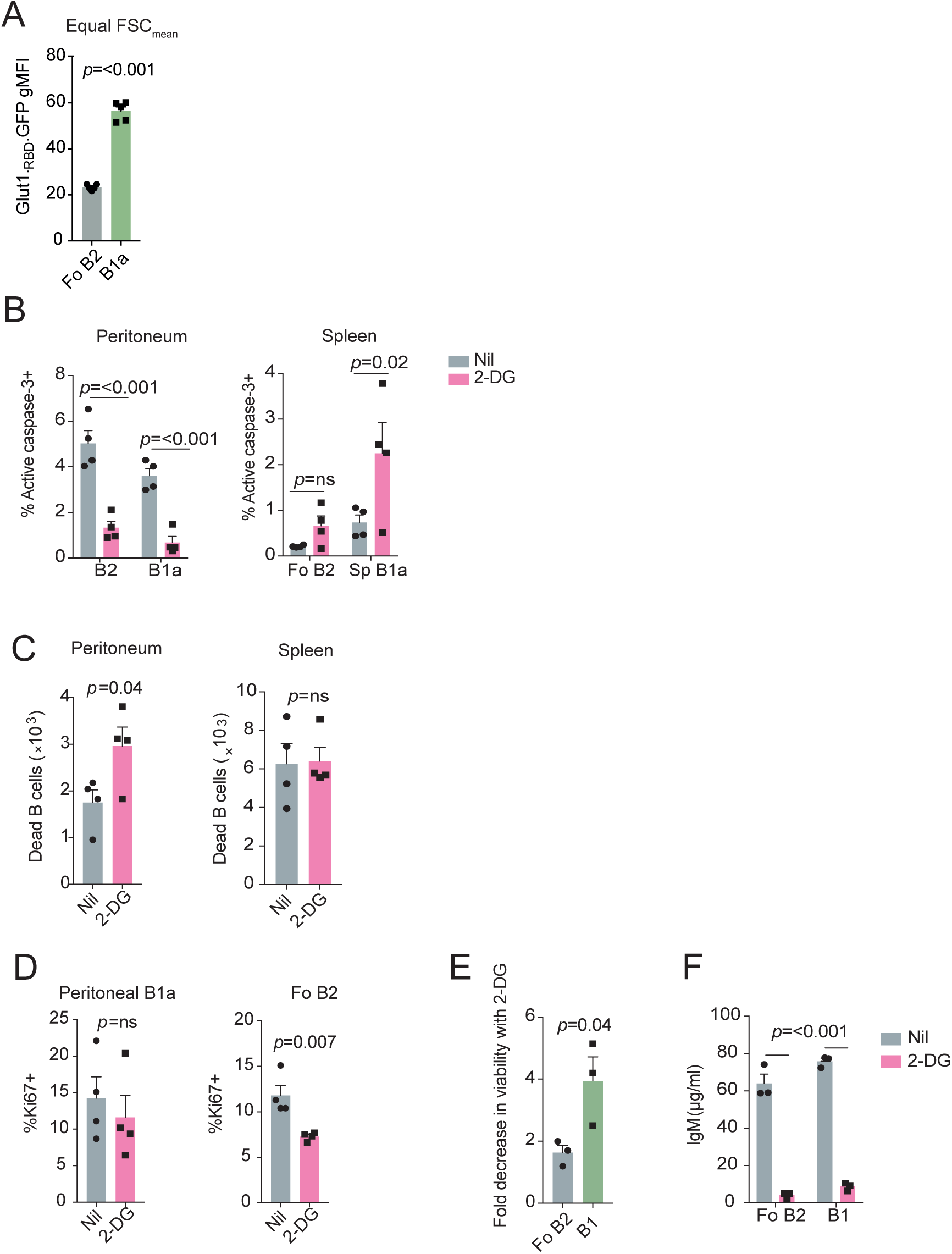
**A. Glut1 expression is higher in B1a B cells than Fo B2 B cells after adjustment for size**. Geometric mean fluorescence of Glut1._RBD_.GFP staining in B1a and B2 cells from the spleen and peritoneum gated so that each population has the same mean flow cytometric forward scatter. Each point represents one C57BL/6 mouse. Mean ±SEM is depicted. Unpaired *t*-test used. Data from Figure 2A. **B. Effect of 2-deoxyglucose on B-cell death *in vivo***. Wild-type C57BL/6 mice were treated with 2-DG as in Figure 2H. Cell death was defined as failure to exclude viability dye, assessed by flow cytometry of cells extracted from the indicated location. Each point represents one mouse. Mean ±SEM is depicted. Unpaired *t*-test used. Representative of two independent experiments. **C. Effect of 2-deoxyglucose on B-cell apoptosis *in vivo***. Wild-type C57BL/6 mice were treated with 2-DG as in Figure 2H. Apoptosis was assessed by intracellular staining for active caspase-3, assessed by flow cytometry of cells extracted from the indicated location. Each point represents one mouse. Mean ±SEM is depicted. Unpaired *t*-test used. Representative of two independent experiments. **D. Effect of 2-deoxyglucose on B-cell proliferation *in vivo***. Wild-type C57BL/6 mice were treated with 2-DG as in Figure 2H. Cell proliferation was assessed by intracellular staining for Ki67, assessed by flow cytometry of cells extracted from the indicated location. Each point represents one mouse. Mean ±SEM is depicted. Unpaired *t*-test used. Representative of two independent experiments. **E. Effect of 2-deoxyglucose on B-cell viability in culture**. Peritoneal B1 (CD19^+^CD23^−^) and splenic follicular B2 B cells (CD19^+^CD23^+^) were isolated by flow cytometry and cultured in the presence or absence of 2.5mM 2-deoxyglucose for 24 hours, following stimulation by 0.5µM ODN1826. Data are presented as fold decrease in viability with treatment, as assessed by measurement of exclusion of viability dye by flow cytometry. Each point represents cells isolated from a pool of 5 C57BL/6 mice. Mean ±SEM is depicted. Unpaired *t*-test used. Representative of two independent experiments. **F. Effect of 2-deoxyglucose on IgM secretion in culture**. IgM production was assessed by ELISA of supernatant from (**F**). Mean ±SEM is depicted. Two-way ANOVA with Sidak correction for multiple comparison used. Representative of two independent experiments.

**Supplementary Figure 2.**
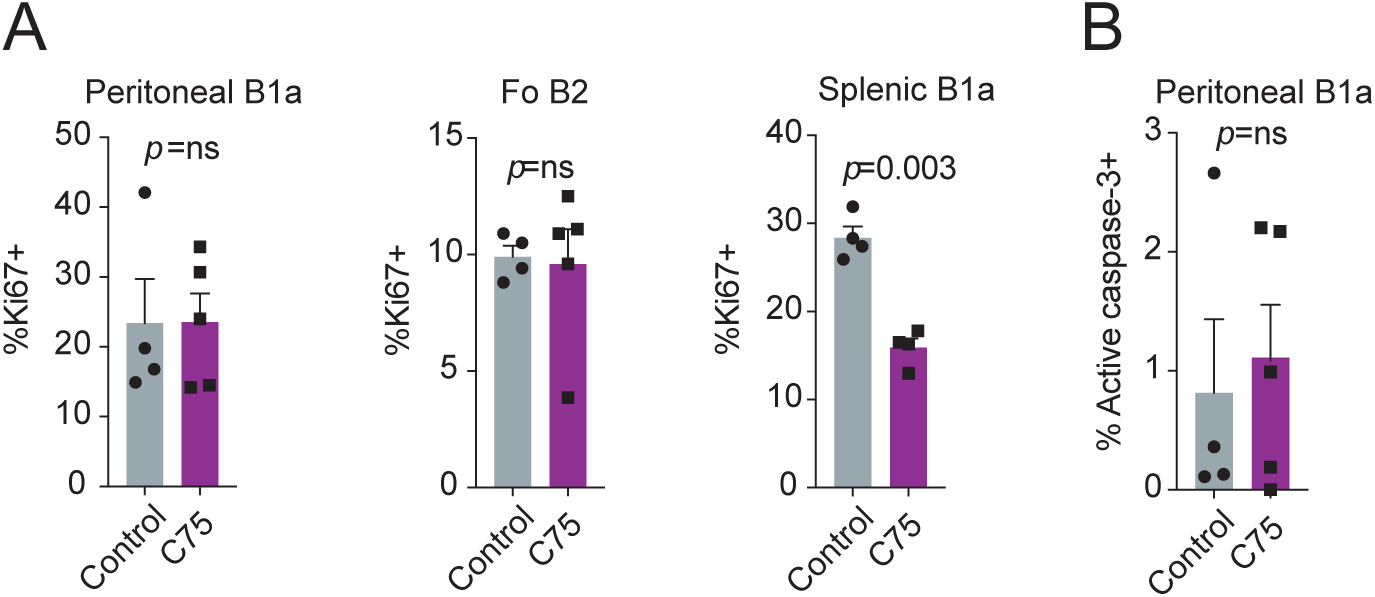
**A. Effect of C75 on B-cell proliferation *in vivo***. Wild-type C57BL/6 mice were treated with C75 as in Figure 3J-K. Cell proliferation was assessed by intracellular staining for Ki67, assessed by flow cytometry of cells extracted from the indicated location. Each point represents one mouse. Mean ±SEM is depicted. Unpaired *t*-test used. Representative of two independent experiments. **B. Effect of C75 on B-cell apoptosis *in vivo***. Wild-type C57BL/6 mice were treated with C75 as in Figure 3J-K. Apoptosis was assessed by intracellular staining for active caspase-3, assessed by flow cytometry of cells extracted from the indicated location. Each point represents one mouse. Mean ±SEM is depicted. Unpaired *t*-test used. Representative of two independent experiments.

**Supplementary Figure 3.**
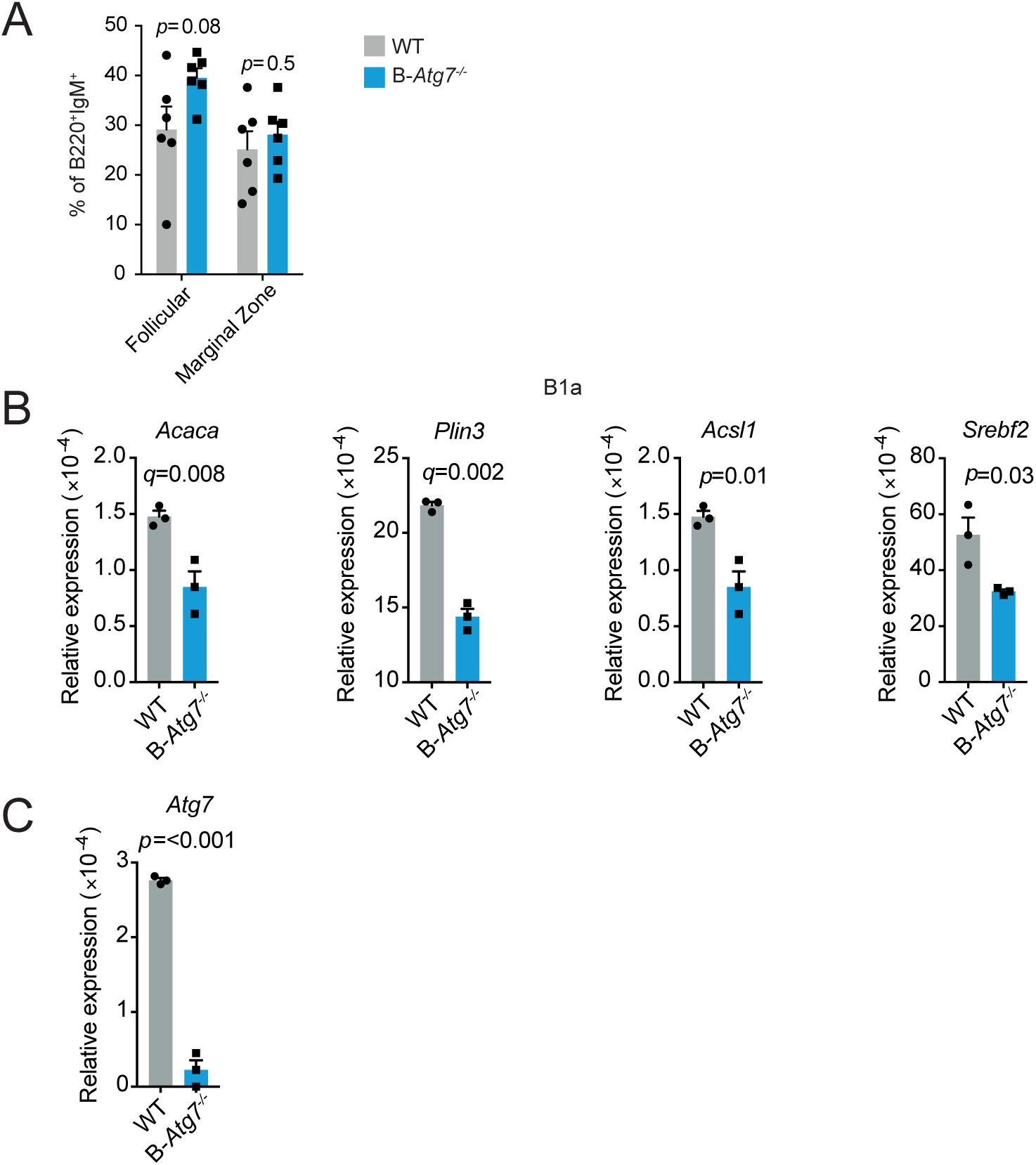
**A. Follicular and marginal B cell populations are intact in B-*Atg7*^−/−^ mice**. Percentages of splenic follicular B2 B cells (CD23^+^) and marginal zone B cells (CD23^lo/−^CD21^hi^) in the B220^+^IgM^+^ population. Each point represents one mouse. Results pooled from two independent experiments. Mean ±SEM is depicted. Two-way ANOVA with Sidak correction for multiple testing used. **B. Loss of autophagy affects metabolic gene expression**. Quantitative RT-qPCR gene expression data for peritoneal B1a B cells from control and B-*Atg7*^−/−^ mice from experiment presented in Figure 5A. Data are relative to β-actin. Each data point is the mean of two technical replicates, representing an individual mouse. Mean ±SEM is depicted. Unpaired *t*-test used. β-values for *Acaca* and *Plin3* are adjusted for multiple testing using FDR method (5% threshold). The post-hoc *p*-values for *Acsl, Srebf2 and Atg7* are presented unadjusted. **C. Mb1-cre efficiently deletes *Atg7* in peritoneal B1a B cells**. Deletion efficiency of Mb1-cre for *Atg7^F/F^* in B1a B cells in control and B-*Atg7*^−/−^ mice from experiment presented in Figure 5A. Data are relative to β-actin. Each data point is the mean of two technical replicates, representing an individual mouse. Mean ±SEM is depicted. Unpaired *t*-test used.

**Supplementary Table 1.**
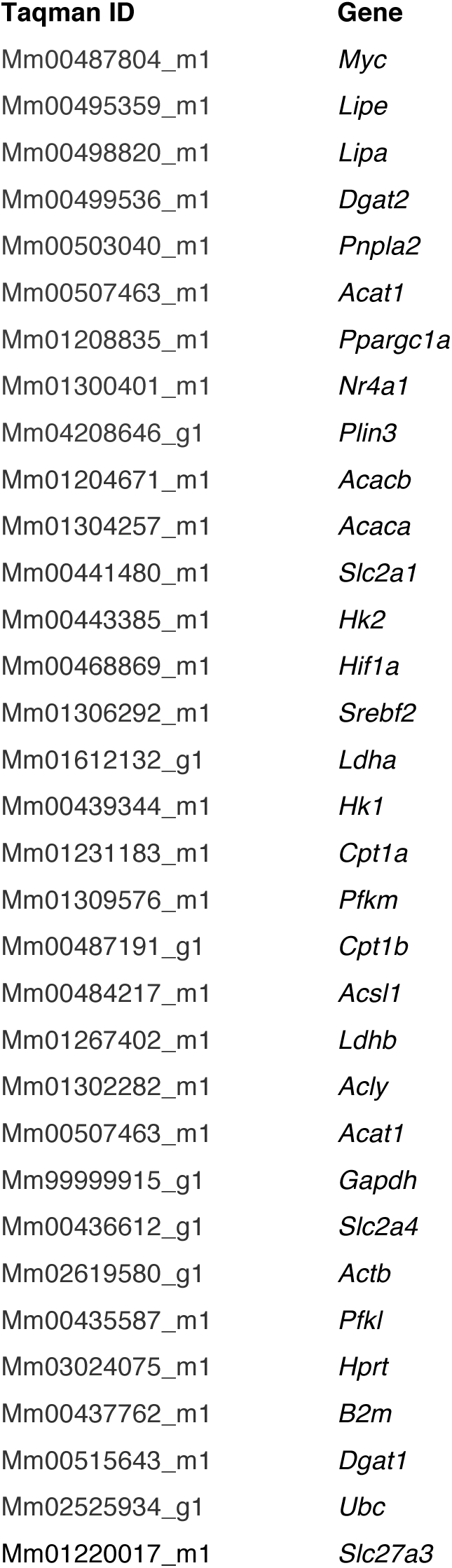
**List of primers used in Biomark experiments and other qPCR**

## References

Ansel, K.M., R.B.S. Harris, and J.G. Cyster. 2002. CXCL13 is required for B1 cell homing, natural antibody production, and body cavity immunity. Immunity. 16: 67–76.

Arnold, J., D. Murera, F. Arbogast, J.-D. Fauny, S. Muller, and F. Gros. 2015. Autophagy is dispensable for B-cell development but essential for humoral autoimmune responses. Cell Death Differ. 1–12. doi:10.1038/cdd.2015.149.

Barber, C.L., E. Montecino-Rodriguez, and K. Dorshkind. 2011. Reduced production of B-1-specified common lymphoid progenitors results in diminished potential of adult marrow to generate B-1 cells. PNAS. 108:13700–13704. doi:10.1073/pnas.1107172108.

Baumgarth, N. 2010. The double life of a B-1 cell: self-reactivity selects for protective effector functions. Nat Rev Immunol. 11:34–46. doi:10.1038/nri2901.

Bhatt, A.P., S.R. Jacobs, A.J. Freemerman, L. Makowski, J.C. Rathmell, D.P. Dittmer, and B. Damania. 2012. Dysregulation of fatty acid synthesis and glycolysis in non-Hodgkin lymphoma. PNAS. 109:11818–11823. doi:10.1073/pnas.1205995109.

Buck, M.D., D. O’Sullivan, and E.L. Pearce. 2015. T cell metabolism drives immunity. Journal of Experimental Medicine. 212:1345–1360. doi:10.1084/jem.20151159.

Buck, M.D., D. O’Sullivan, R.I.K. Geltink, J.D. Curtis, C.-H. Chang, D.E. Sanin, J. Qiu, O. Kretz, D. Braas, G.J.W. van der Windt, Q. Chen, S.C.-C. Huang, C.M. O'Neill, B.T. Edelson, E.J. Pearce, H. Sesaki, T.B. Huber, A.S. Rambold, and E.L. Pearce. 2016. Mitochondrial Dynamics Controls T Cell Fate through Metabolic Programming. Cell. 166:63–76. doi:10.1016/j.cell.2016.05.035.

Caro-Maldonado, A., R. Wang, A.G. Nichols, M. Kuraoka, S. Milasta, L.D. Sun, A.L. Gavin, E.D. Abel, G. Kelsoe, D.R. Green, and J.C. Rathmell. 2014. Metabolic Reprogramming Is Required for Antibody Production That Is Suppressed in Anergic but Exaggerated in Chronically BAFF-Exposed B Cells. The Journal of Immunology. 192:3626–3636. doi: 10.4049/jimmunol.1302062.

Chen, M., M.J. Hong, H. Sun, L. Wang, X. Shi, B.E. Gilbert, D.B. Corry, F. Kheradmand, and J. Wang. 2014. Essential role for autophagy in the maintenance of immunological memory against influenza infection. Nature Medicine. 20:503–510. doi:10.1038/nm.3521.

Chen, Y., Y.B. Park, E. Patel, and G.J. Silverman. 2009. IgM Antibodies to Apoptosis-Associated Determinants Recruit C1q and Enhance Dendritic Cell Phagocytosis of Apoptotic Cells. J Immunol. 182:6031–6043. doi:10.4049/jimmunol.0804191.

Clarke, A.J., U. Ellinghaus, A. Cortini, A. Stranks, A.K. Simon, M. Botto, and T.J. Vyse. 2014. Autophagy is activated in systemic lupus erythematosus and required for plasmablast development. Annals of the Rheumatic Diseases. doi:10.1136/annrheumdis-2013-204343.

Doughty, C.A. 2006. Antigen receptor-mediated changes in glucose metabolism in B lymphocytes: role of phosphatidylinositol 3-kinase signaling in the glycolytic control of growth. Blood. 107:4458–4465. doi:10.1182/blood-2005-12-4788.

Eng, K.E., M.D. Panas, G.B. Karlsson Hedestam, and G.M. McInerney. 2010. A novel quantitative flow cytometry-based assay for autophagy. Autophagy. 6:634–641. doi:10.4161/auto.6.5.12112.

Esplin, B.L., R.S. Welner, Q. Zhang, L.A. Borghesi, and P.W. Kincade. 2009. A differentiation pathway for B1 cells in adult bone marrow. PNAS. 106:5773–5778. doi:10.1073/pnas.0811632106.

García-Prat, L., M. Martinez-Vicente, E. Perdiguero, L. Ortet, J. Rodríguez-Ubreva, E. Rebollo, V. Ruiz-Bonilla, S. Gutarra, E. Ballestar, A.L. Serrano, M. Sandri, and P. Muñoz-Cánoves. 2016. Autophagy maintains stemness by preventing senescence. Nature. 529:37–42. doi:10.1038/nature16187.

Guan, J.-L., A.K. Simon, M. Prescott, J.A. Menendez, F. Liu, F. Wang, C. Wang, E. Wolvetang, A. Vazquez-Martin, and J. Zhang. 2013. Autophagy in stem cells. Autophagy. 9:830–849. doi:10.4161/auto.24132.

Hauck, A.K., and D.A. Bernlohr. 2016. Oxidative stress and lipotoxicity. J. Lipid Res. 57:1976–1986. doi:10.1194/jlr.R066597.

Hayakawa, K., A.M. Formica, J. Brill-Dashoff, S.A. Shinton, D. Ichikawa, Y. Zhou, H.C. Morse III, and R.R. Hardy. 2016. Early generated B1 B cells with restricted BCRs become chronic lymphocytic leukemia with continued c-Myc and low Bmf expression. J. Exp. Med. 213:3007–3024. doi:10.1084/jem.20160712.

Hayakawa, K., R.R. Hardy, A.M. Stall, and L.A. Herzenberg. 1986. Immunoglobulin-bearing B cells reconstitute and maintain the murine Ly-1 B cell lineage. Eur. J. Immunol. 16:1313–1316. doi:10.1002/eji.1830161021.

Hobeika, E., S. Thiemann, B. Storch, H. Jumaa, P.J. Nielsen, R. Pelanda, and M. Reth. 2006. Testing gene function early in the B cell lineage in mb1-cre mice. Proc. Natl. Acad. Sci. U.S.A. 103:13789–13794. doi:10.1073/pnas.0605944103.

Jitschin, R., A.D. Hofmann, H. Bruns, A. Giessl, J. Bricks, J. Berger, D. Saul, M.J. Eckart, A. Mackensen, and D. Mougiakakos. 2014. Mitochondrial metabolism contributes to oxidative stress and reveals therapeutic targets in chronic lymphocytic leukemia. Blood. 123:2663–2672. doi:10.1182/blood-2013-10-532200.

Kabat, A.M., O.J. Harrison, and T. Riffelmacher. 2016. The autophagy gene Atg16l1 differentially regulates Treg and TH2 cells to control intestinal inflammation. eLife. doi: 10.7554/eLife.12444.001.

Kaminski, D.A., and J. Stavnezer. 2006. Enhanced IgA Class Switching in Marginal Zone and B1 B Cells Relative to Follicular/B2 B Cells. J Immunol. 177:6025–6029. doi:10.4049/j immunol.177.9.6025.

Kaur, J., and J. Debnath. 2015. Autophagy at the crossroads of catabolism and anabolism. Nat Rev Mol Cell Biol. 16:461–472. doi:10.1038/nrm4024.

Khuda, S.E., W.M. Loo, S. Janz, B. Van Ness, and L.D. Erickson. 2008. Deregulation of c-Myc Confers Distinct Survival Requirements for Memory B Cells, Plasma Cells, and Their Progenitors. J Immunol. 181:7537–7549. doi:10.4049/jimmunol.181.11.7537.

Krop, I., A.R. de Fougerolles, R.R. Hardy, M. Allison, M.S. Schlissel, and D.T. Fearon. 1996. Self-renewal of B-1 lymphocytes is dependent on CD19. Eur. J. Immunol. 26:238–242. doi:10.1002/eji.1830260137.

Levine, B., N. Mizushima, and H.W. Virgin. 2011. Autophagy in immunity and inflammation. Nature. 469:323–335. doi:10.1038/nature09782.

Liu, K., and M.J. Czaja. 2012. Regulation of lipid stores and metabolism by lipophagy. Cell Death Differ. 20:3–11. doi:10.1038/cdd.2012.63.

Loftus, T.M., D.E. Jaworsky, G.L. Frehywot, C.A. Townsend, G.V. Ronnett, M.D. Lane, and F. P. Kuhajda. 2000. Reduced food intake and body weight in mice treated with fatty acid synthase inhibitors. Science. 288: 2379–2381.

Macintyre, A.N., V.A. Gerriets, A.G. Nichols, R.D. Michalek, M.C. Rudolph, D. Deoliveira, S.M. Anderson, E.D. Abel, B.J. Chen, L.P. Hale, and J.C. Rathmell. 2014. The Glucose Transporter Glut1 Is Selectively Essential for CD4 T Cell Activation and Effector Function. Cell Metabolism. 20:61–72. doi:10.1016/j.cmet.2014.05.004.

Manel, N., F.J. Kim, S. Kinet, N. Taylor, M. Sitbon, and J.-L. Battini. 2003. The ubiquitous glucose transporter GLUT-1 is a receptor for HTLV. Cell. 115: 449–459.

Michalek, R.D., V.A. Gerriets, S.R. Jacobs, A.N. Macintyre, N.J. MacIver, E.F. Mason, S.A. Sullivan, A.G. Nichols, and J.C. Rathmell. 2011. Cutting Edge: Distinct Glycolytic and Lipid Oxidative Metabolic Programs Are Essential for Effector and Regulatory CD4+ T Cell Subsets. The Journal of Immunology. 186:3299–3303. doi: 10.4049/jimmunol.1003613.

Miller, B.C., Z. Zhao, L.M. Stephenson, K. Cadwell, H.H. Pua, H.K. Lee, N.N. Mizushima, A. Iwasaki, Y.-W. He, W. Swat, and H.W. Virgin. 2008. The autophagy gene ATG5 plays an essential role in B lymphocyte development. Autophagy. 4: 309–314.

Montecino-Rodriguez, E., and K. Dorshkind. 2012. B-1 B Cell Development in the Fetus and Adult. Immunity. 36:13–21. doi:10.1016/j.immuni.2011.11.017.

Mortensen, M., E.J. Soilleux, G. Djordjevic, R. Tripp, M. Lutteropp, E. Sadighi-Akha, A.J. Stranks, J. Glanville, S. Knight, S.E. W Jacobsen, K.R. Kranc, and A.K. Simon. 2011. The autophagy protein Atg7 is essential for hematopoietic stem cell maintenance. Journal of Experimental Medicine. 208:455–467. doi:10.1084/jem.20101145.

Nagaraj, R., M.S. Sharpley, F. Chi, D. Braas, Y. Zhou, R. Kim, A.T. Clark, and U. Banerjee. 2017. Nuclear Localization of Mitochondrial TCA Cycle Enzymes as a Critical Step in Mammalian Zygotic Genome Activation. Cell. 168:210–223.e11. doi: 10.1016/j.cell.2016.12.026.

Pan, H., N. Cai, M. Li, G.-H. Liu, and J.C. Izpisua Belmonte. 2013. Autophagic control of cell “stemness” EMBO Mol Med. 5:327–331. doi:10.1002/emmm.201201999.

Pan, Y., T. Tian, C.O. Park, S.Y. Lofftus, S. Mei, X. Liu, C. Luo, J.T. O’Malley, A. Gehad, J.E. Teague, S.J. Divito, R. Fuhlbrigge, P. Puigserver, J.G. Krueger, G.S. Hotamisligil, R.A. Clark, and T.S. Kupper. 2017. Survival of tissue-resident memory T cells requires exogenous lipid uptake and metabolism. Nature. 543:252–256. doi:10.1038/nature21379.

Pearce, E.L., and E.J. Pearce. 2013. Metabolic Pathways in Immune Cell Activation and Quiescence. Immunity. 38:633–643. doi:10.1016/j.immuni.2013.04.005.

Pearce, E.L., M.C. Walsh, P.J. Cejas, G.M. Harms, H. Shen, L.-S. Wang, R.G. Jones, and Y. Choi. 2009. Enhancing CD8 T-cell memory by modulating fatty acid metabolism. Nature. 460:103–107. doi:10.1038/nature08097.

Pengo, N., M. Scolari, L. Oliva, E. Milan, F. Mainoldi, A. Raimondi, C. Fagioli, A. Merlini, E. Mariani, E. Pasqualetto, U. Orfanelli, M. Ponzoni, R. Sitia, S. Casola, and S. Cenci. 2013. Plasma cells require autophagy for sustainable immunoglobulin production. Nature Immunology. 14:298–305. doi:10.1038/ni.2524.

Puleston, D.J., H. Zhang, T.J. Powell, E. Lipina, S. Sims, I. Panse, A.S. Watson, V. Cerundolo, A.R. Townsend, P. Klenerman, and A.K. Simon. 2014. Autophagy is a critical regulator of memory CD8 +T cell formation. eLife. 3:2516–21. doi:10.7554/eLife.03706.

Rambold, A.S., S. Cohen, and J. Lippincott-Schwartz. 2015. Fatty acid trafficking in starved cells: regulation by lipid droplet lipolysis, autophagy, and mitochondrial fusion dynamics. Dev. Cell. 32:678–692. doi:10.1016/j.devcel.2015.01.029.

Riffelmacher, T., A. Clarke, F.C. Richter, A. Stranks, S. Pandey, S. Danielli, P. Hublitz, Z. Yu, E. Johnson, T. Schwerd, J. McCullagh, H. Uhlig, S.E.W. Jacobsen, and A.K. Simon. 2017. Autophagy-Dependent Generation of Free Fatty Acids Is Critical for Normal Neutrophil Differentiation. Immunity. 47:466–480.e5. doi:10.1016/j.immuni.2017.08.005.

Rozovski, U., I. Hazan-Halevy, M. Barzilai, M.J. Keating, and Z. Estrov. 2016. Metabolism pathways in chronic lymphocytic leukemia. Leukemia & Lymphoma. 57:758–765. doi:10.3109/10428194.2015.1106533.

Singh, R., S. Kaushik, Y. Wang, Y. Xiang, I. Novak, M. Komatsu, K. Tanaka, A.M. Cuervo, and M.J. Czaja. 2009. nature07976. Nature. 458:1131–1135. doi:10.1038/nature07976.

Sukumar, M., J. Liu, G.U. Mehta, S.J. Patel, R. Roychoudhuri, J.G. Crompton, C.A. Klebanoff, Y. Ji, P. Li, Z. Yu, G.D. Whitehill, D. Clever, R.L. Eil, D.C. Palmer, S. Mitra, M. Rao, K. Keyvanfar, D.S. Schrump, E. Wang, F.M. Marincola, L. Gattinoni, W.J. Leonard, P. Muranski, T. Finkel, and N.P. Restifo. 2016. Mitochondrial Membrane Potential Identifies Cells with Enhanced Stemness for Cellular Therapy. Cell Metabolism. 23:63–76. doi:10.1016/j.cmet.2015.11.002.

Sukumar, M., J. Liu, Y. Ji, M. Subramanian, J.G. Crompton, Z. Yu, R. Roychoudhuri, D.C. Palmer, P. Muranski, E.D. Karoly, R.P. Mohney, C.A. Klebanoff, A. Lal, T. Finkel, N.P. Restifo, and L. Gattinoni. 2013. Inhibiting glycolytic metabolism enhances CD8+ T cell memory and antitumor function. J. Clin. Invest. 123:4479–4488. doi:10.1172/JCI69589.

Tehranchi, R., P.S. Woll, K. Anderson, N. Buza-Vidas, T. Mizukami, A.J. Mead, I. Astrand-Grundström, B. Strömbeck, A. Horvat, H. Ferry, R.S. Dhanda, R. Hast, T. Rydén, P. Vyas, G. Göhring, B. Schlegelberger, B. Johansson, E. Hellström-Lindberg, A. List, L. Nilsson, and S.E.W. Jacobsen. 2010. Persistent malignant stem cells in del(5q) myelodysplasia in remission. N. Engl. J. Med. 363:1025–1037. doi: 10.1056/NEJMoa0912228.

Tili, E., J.J. Michaille, Z. Luo, S. Volinia, L.Z. Rassenti, T.J. Kipps, and C.M. Croce. 2012. The down-regulation of miR-125b in chronic lymphocytic leukemias leads to metabolic adaptation of cells to a transformed state. Blood. 120:2631–2638. doi:10.1182/blood-2012-03-415737.

Tung, J.W., M.D. Mrazek, Y. Yang, L.A. Herzenberg, and L.A. Herzenberg. 2006. Phenotypically distinct B cell development pathways map to the three B cell lineages in the mouse. Proc. Natl. Acad. Sci. U.S.A. 103:6293–6298. doi:10.1073/pnas.0511305103.

Vannini, N., M. Girotra, O. Naveiras, G. Nikitin, V. Campos, S. Giger, A. Roch, J. Auwerx, and M.P. Lutolf. 2016. Specification of haematopoietic stem cell fate via modulation of mitochondrial activity. Nat Commun. 7:1–9. doi:10.1038/ncomms13125.

Watson, A.S., T. Riffelmacher, A. Stranks, O. Williams, J. De Boer, K. Cain, M. MacFarlane, J. McGouran, B. Kessler, S. Khandwala, O. Chowdhury, D. Puleston, K. Phadwal, M. Mortensen, D. Ferguson, E. Soilleux, P. Woll, S. Jacobsen, and A.K. Simon. 2015. Autophagy limits proliferation and glycolytic metabolism in acute myeloid leukemia. Cell Death Discovery. 1:15008–10. doi:10.1038/cddiscovery.2015.8.

Wilfling, F., J.T. Haas, T.C. Walther, and R.V. Farese Jr. 2014. ScienceDirectLipid droplet biogenesis. Current Opinion in Cell Biology. 29:39–45. doi:10.1016/j.ceb.2014.03.008.

Wilhelm, C., O.J. Harrison, V. Schmitt, M. Pelletier, S.P. Spencer, J.F. Urban Jr., M. Ploch, T.R. Ramalingam, R.M. Siegel, and Y. Belkaid. 2016. Critical role of fatty acid metabolism in ILC2-mediated barrier protection during malnutrition and helminth infection. J. Exp. Med. 213:1409–1418. doi:10.1084/jem.20151448.

Yang, Y., J.W. Tung, E.E.B. Ghosn, L.A. Herzenberg, and L.A. Herzenberg. 2007. Division and differentiation of natural antibody-producing cells in mouse spleen. Proc. Natl. Acad. Sci. U.S.A. 104:4542–4546. doi:10.1073/pnas.0700001104.

Youle, R.J., and A.M. van der Bliek. 2012. Mitochondrial Fission, Fusion, and Stress. Science. 337:1062–1065. doi:10.1126/science.1219855.

